# Thrombospondin 2 is a key determinant of fibrogenesis in NAFLD

**DOI:** 10.1101/2023.06.01.543250

**Authors:** Takefumi Kimura, Takanobu Iwadare, Shun-ichi Wakabayashi, Seema Kuldeep, Tomoyuki Nakajima, Tomoo Yamazaki, Daiki Aomura, Hamim Zafar, Mai Iwaya, Takeshi Uehara, Sai P Pydi, Naoki Tanaka, Takeji Umemura

## Abstract

Hepatic overexpression of the thrombospondin 2 gene (THBS2) and elevated levels of circulating thrombospondin 2 (TSP2) have been observed in patients with chronic liver disease. The current study aimed to identify the specific cells expressing THBS2/TSP2 in non-alcoholic fatty liver disease (NAFLD) and investigate the underlying mechanism behind THBS2/TSP2 up-regulation. Comprehensive NAFLD liver gene datasets, including single-cell RNA sequencing (scRNA-seq), in-house NAFLD liver tissue, and LX-2 cells derived from human hepatic stellate cells (HSCs), were analyzed using a combination of computational biology, genetic, immunological, and pharmacological approaches. Analysis of the genetic dataset revealed the presence of 1433 variable genes in patients with advanced fibrosis NAFLD, with THBS2 ranked among the top 2 genes. Quantitative polymerase chain reaction examination of NAFLD livers showed a significant correlation between THBS2 expression and fibrosis stage (r=0.349, p<0.001). In support of this, scRNA-seq data and in situ hybridization demonstrated that the THBS2 gene was highly expressed in HSCs of NAFLD patients with advanced fibrosis. Pathway analysis of the gene dataset revealed THBS2 expression to be associated with the transforming growth factor beta (TGFβ) pathway and collagen gene activation. Moreover, the activation of LX-2 cells with TGFβ increased THBS2/TSP2 and collagen expression independently of the TGFβ-SMAD2/3 pathway. THBS2 gene knockdown significantly decreased collagen expression in LX-2 cells. In conclusion, THBS2/TSP2 is highly expressed in HSCs and plays a role in regulating fibrogenesis in NAFLD patients. THBS2/TSP2 may therefore represent a potential target for anti-fibrotic therapy in NAFLD. (241 words)

**One-sentence summaries:** Thrombospondin 2 represent a potential target for anti-fibrotic therapy in NAFLD.

## Introduction

Non-alcoholic fatty liver disease (NAFLD) is a global health concern (*1, 2*) that encompasses a spectrum of conditions, ranging from non-alcoholic fatty liver to non-alcoholic steatohepatitis (NASH) (*3*). NASH is characterized by the accumulation of fat in the liver accompanied by inflammation and scarring, which can lead to cirrhosis and liver cancer (*4*). Several large clinical trials have suggested that the degree of liver fibrosis is closely related to prognosis in NAFLD (*5–7*). However, the mechanisms underlying the progression of liver fibrosis are not fully understood, and effective therapeutic strategies for NAFLD have yet to be established (*8, 9*).

Thrombospondins are a group of proteins that are produced and secreted by various cells (*10, 11*). The glycoproteins are characterized by multiple domains and exhibit calcium-binding capability (*12*). Thrombospondins interact with a wide range of substances in the body, including cytokines, growth factors, receptors, and components of the extracellular matrix (ECM) (*13*). Of the five different types of thrombospondins, thrombospondin 2 (TSP2) stands out as a particularly unique member. TSP2 is encoded by the thrombospondin 2 (THBS2) gene and is involved in such processes as fibrin formation, bone growth, maintenance of normal blood vessel density, blood clotting, and cell adhesion (*14*). In the skin, TSP2 is mainly produced by fibroblasts and smooth muscle cells and is believed to play a role in the process of wound repair and tissue remodeling (*14*). Recently, several studies have reported that THBS2 is upregulated in the fibrotic liver in NAFLD, with secreted TSP2 emerging as a potential biomarker for disease progression (*15, 16*). These results were supported by a large investigation of NAFLD patients with diabetes mellitus or metabolic-associated fatty liver disease (*17, 18*). Furthermore, a correlation between the degree of fibrosis and inflammation and serum TSP2 levels was observed not only in NAFLD, but also in hepatitis C virus infected patients (*19, 20*). Despite these advances, however, the specific liver cell populations responsible for THBS2/TSP2 expression in NAFLD, the underlying mechanisms governing expression dynamics, and the precise role of THBS2/TSP2 in the pathogenesis of NAFLD remain largely unknown.

To address the above issues, we used comprehensive genetic data, single-cell RNA sequencing (scRNA-seq) datasets, in situ hybridization data, and in vitro models to identify THBS2/TSP2-expressing cells in NAFLD, elucidate their expression mechanisms, and investigate their potential therapeutic applications. We go on to show that THBS2/TSP2 is highly expressed in hepatic stellate cells (HSCs) and plays a role in regulating the fibrotic process in patients with NAFLD.

## Results

### Hepatic THBS2 expression is markedly increased in NAFLD patients with advanced fibrosis

We first analyzed a RNA sequencing dataset (GSE49541) to determine the extent to which THBS2 was highly expressed in the liver of patients with NAFLD (*21, 22*). When comparing mild fibrosis (F0-1) with advanced fibrosis (F3-4), 1433 out of 53242 genes showed significant variation (Figure 1a). Characteristically, THBS2 ranked second of the 1433 variable genes exhibited by advanced fibrosis NAFLD (Figure 1b). The Log2 fold change for THBS2 was - 2.113, and -log10 (P value) was 15.821. The expression values of THBS2 in mild (F0-1) and advanced (F3-4) fibrosis cases are shown in Figure 1c. While THBS2 expression was not high for mild fibrosis, elevated THBS2 expression was evident in many advanced fibrosis cases. qPCR using 96 biopsied NAFLD samples also revealed a correlation between fibrosis stage and hepatic THBS2 expression (r=0.349, P=0.00046) (Figure 1d), thus validating the RNA sequencing data (Figure 1b, c). Similarly, THBS2 expression was significantly higher in patients with greater fibrosis in comparisons of F0-1 vs. F2-4 and F0-2 vs. F3-4 (P=0.0015 and P<0.0001, respectively) (Figure 1e).

**Fig. 1.**
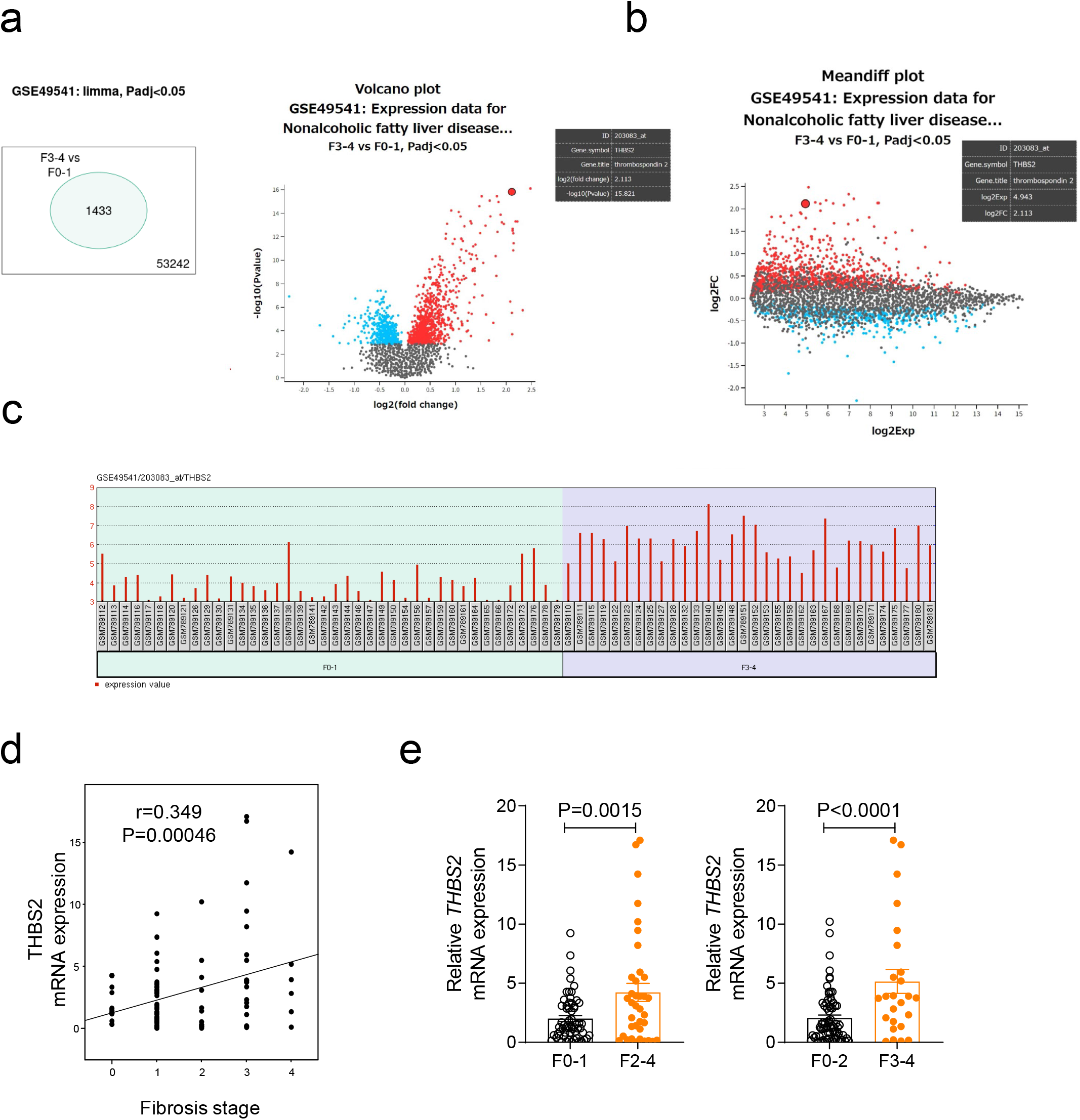
THBS2 gene expression is upregulated in NAFLD patients with advanced fibrosis. **a-c,** Gene expression analysis using a NAFLD liver-derived microarray dataset (GSE49541). Number of genes **(a, left)**, volcano plot **(a, right)**, and mean differentiation plot **(b)** with an adjusted P value of <0.05 when comparing F3-4 cases with F0-1 cases. The large red dot indicates THBS2. **(c)** THBS2 expression levels by NAFLD case in GSE49541 (F0-1, n=40; F3-4, n=32). **d, e,** Relative THBS2 mRNA expression levels in NAFLD liver tissue samples from our institution. Relative THBS2 mRNA expression levels (n=96) by fibrosis stage **(d)**, F0-1 (n=59) vs. F2-4 (n=37) and F0-2 (n=71) vs. F3-4 (n=25) **(e)**. Data are presented as the mean ± standard error of the mean. Correlation analysis was conducted by Spearman’s test **(d)** or the two-tailed Student’s *t*-test **(e)**.

### Enrichment analysis of comprehensive genetic data: association of THBS2 with collagen-related genes and TGFβ pathway

Enrichment analysis was performed using Metascape (*23*) using the same gene dataset (GSE49541) as in the previous section (*21, 22*). In NAFLD with advanced fibrosis, the TGFβ-signaling pathway was activated in addition to the organization of collagen-containing NABA core matrisome and ECM (Figure 2a, underlined).

**Fig. 2.**
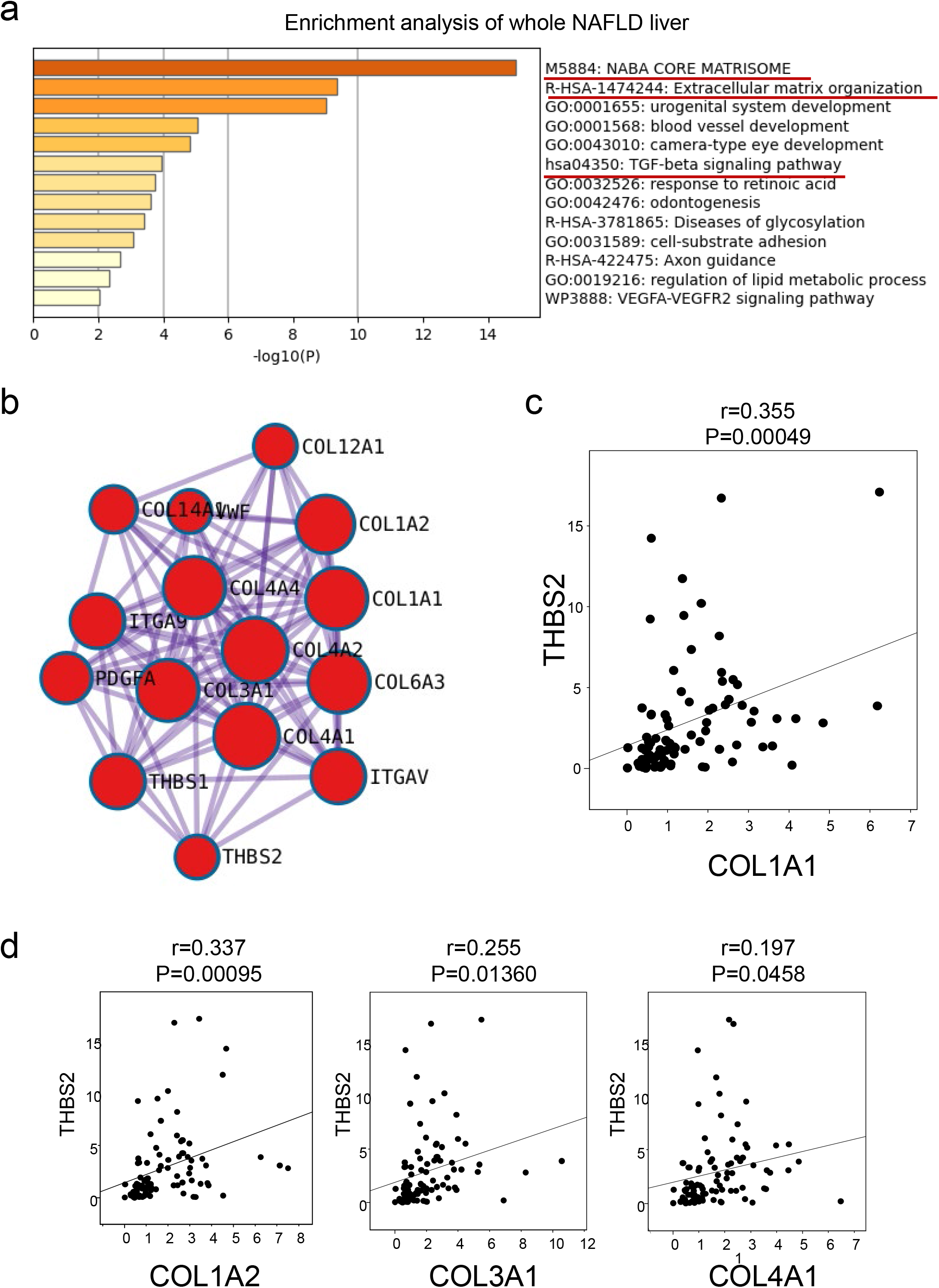
Hepatic THBS2 is associated with collagen genes in NAFLD. **a, b**, Enrichment analysis **(a)** and network analysis **(b)** in a NAFLD liver gene set (GSE49541) using Metascape (*23*). **(b)** Group of genes showing strong linkage to THBS2. This is a partial expansion of Supp. Fig. 1. **c, d,** Correlation of relative mRNA expression levels of THBS2 and COL1A1 **(c)**, as well as of THBS2 with COL1A2, COL3A1, and COL4A1 **(d)** in NAFLD liver tissue samples from our institution (n=96). Correlation analysis was conducted by Spearman’s test **(c)**.

Supplementary Figure 1 shows the THBS2 gene network in the progression of fibrotic NAFLD, with Figure 2b providing partial enlargement of the THBS2 nearside. The established MCODE algorithm of Metascape was employed to find the densely connected protein neighborhoods in the network, showing the biological role of each component. Network analysis revealed THBS2 to be associated with collagen-related genes, including COL1A1, COL1A2, COL3A1, and COL4A1. Supportive qPCR data from liver biopsy tissue from our own cohort also showed a significant correlation between THBS2 and collagen-related genes (vs. COL1A1: r=0.355, P=0.00049; vs. COL1A2: r=0.337, P=0.00095; vs. COL3A1: r=0.225, P=0.01360; vs. COL4A1: r=0.197, P=0.0458) (Figure 2c, d). The clinical data of the patients used for qPCR are shown in Table 1.

**Table 1.** Clinico-pathological background of patients from own institution used for qPCR studies.

|  | All (n=96) |
| --- | --- |
|  | Median (IQR) / n (%) |
| Age (years) | 56 (37-65) |
| Male | 50 (52%) |
| Body mass index (kg/m<sup>2</sup>) | 26.1 (24-30) |
| **Laboratory data** |  |
| Albumin (mg/dL) | 4.6 (4.3-4.8) |
| Bilirubin (mg/dL) | 0.91 (0.71-1.18) |
| AST (U/L) | 44 (29-64) |
| ALT (U/L) | 62 (19-273) |
| GGTP (IU/L) | 50 (36-86) |
| Cholinesterase (U/L) | 375 (298-433) |
| Fasting glucose (mg/dL) | 106 (95-116) |
| Insulin (μU/mL) | 13.5 (8.9-26) |
| HOMA-IR | 3.7 (2.4-7.6) |
| Hemoglobin A1c (%) | 5.6 (5.2-6.0) |
| Total cholesterol (mg/dL) | 209 (179-237) |
| LDL cholesterol (mg/dL) | 130 (118-154) |
| HDL cholesterol (mg/dL) | 49 (43-57) |
| Triglycerides (mg/dL) | 134 (97-176) |
| Alpha fetoprotein (ng/mL) | 2.6 (1.9-3.6) |
| Platelets (×10<sup>4</sup>/uL) | 20.4 (16.0-25.6) |
| APRI | 0.67 (0.46-1.28) |
| FIB-4 | 1.52 (0.76-2.5) |
| **Pathology** |  |
| NAFLD activity score | 5 (3-5) |
| Fibrosis stage (0/1/2/3/4) | 15/44/12/19/6 |

### scRNA-seq data and in situ hybridization analysis of fibrotic NAFLD cases confirm the overexpression of THBS2 in HSCs

scRNA-seq data obtained from the GSE174748 and GSE189175 datasets were analyzed using Seurat 4.5 to determine the THBS2-expressing cells in healthy individuals and patients with NAFLD (Figure 3a) (*24, 25*). The THBS2 gene was particularly enriched in HSC populations over other cell types (Figure 3b). Notably, we observed a significant upregulation of THBS2 expression in fibrotic NAFLD patients, particularly in HSCs, when compared with healthy controls and subjects with NAFLD but no fibrosis (Figure 3c). These findings suggested that THBS2 might play a role in the development of fibrosis in NAFLD.

**Fig. 3.**
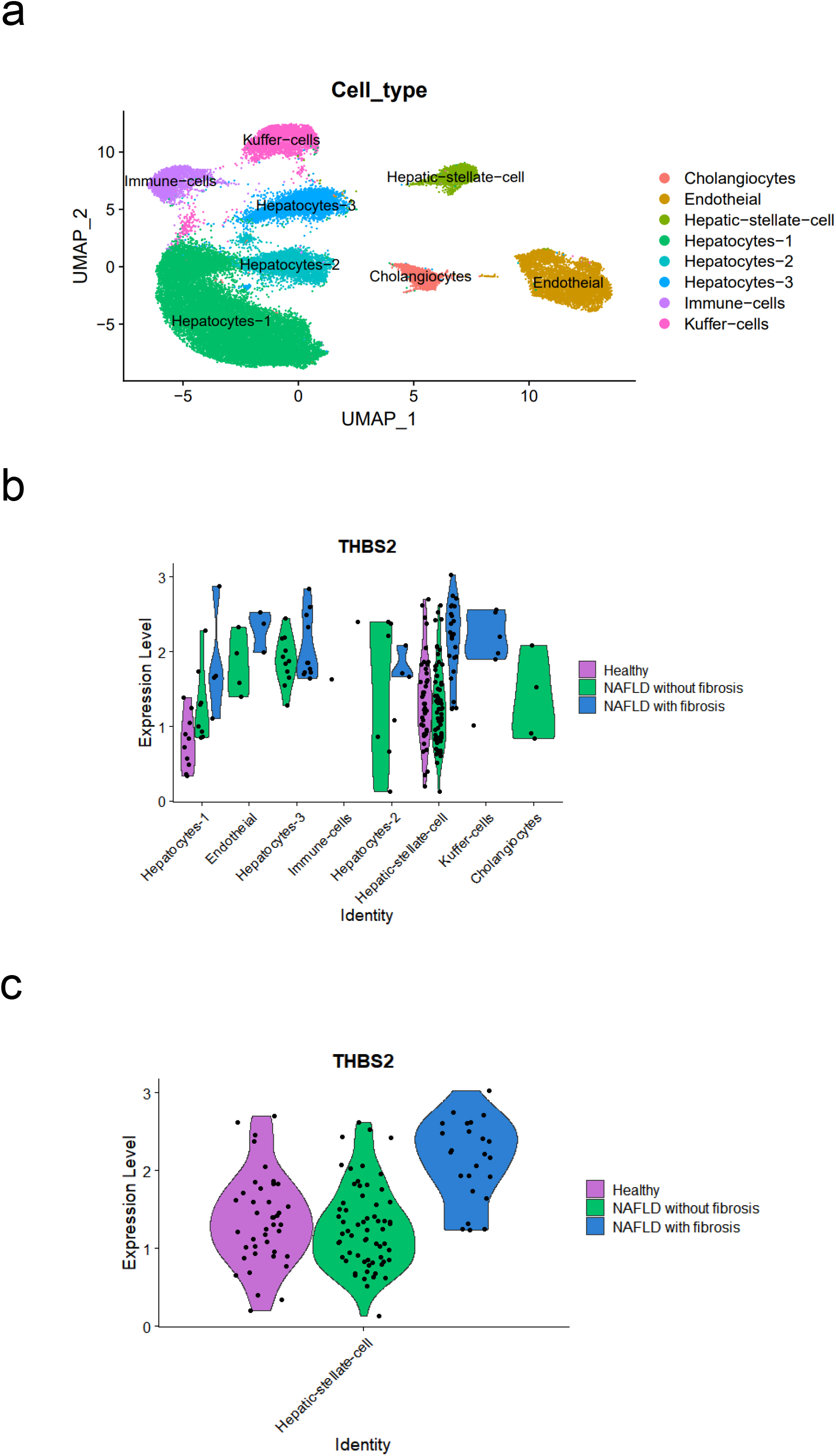
Predominant expression of THBS2 in HSCs by scRNA-seq data analysis. **a-c,** GSE174748 and GSE189175 scRNA-seq datasets were used in this study. See Methods for details. **(a)** Liver cell clusters in the whole dataset. **b, c,** THBS2 expression levels in each cell cluster **(b)** and in HSCs **(c)**. THBS2 expression levels of HSCs are shown in three groups: Healthy, NAFLD with fibrosis, and NAFLD without fibrosis.

To further confirm the expression of THBS2 in specific cell types, we performed in situ hybridization analysis using a THBS2 mRNA probe of liver tissue sections from severely fibrotic NAFLD cases, which revealed positive brown signals indicating THBS2 expression in HSCs in F3 NAFLD livers (Figure 4a-c, left). In contrast, no THBS2 signals were observed in control livers (Figure 4d, left). Upon closer examination (Figure 4a), THBS2 signaling was mainly present around hepatocytes and collagen fibers. These results supported the scRNA-seq data, confirming that HSCs contributed to the elevated expression of THBS2 in fibrotic NAFLD livers.

**Fig. 4.**
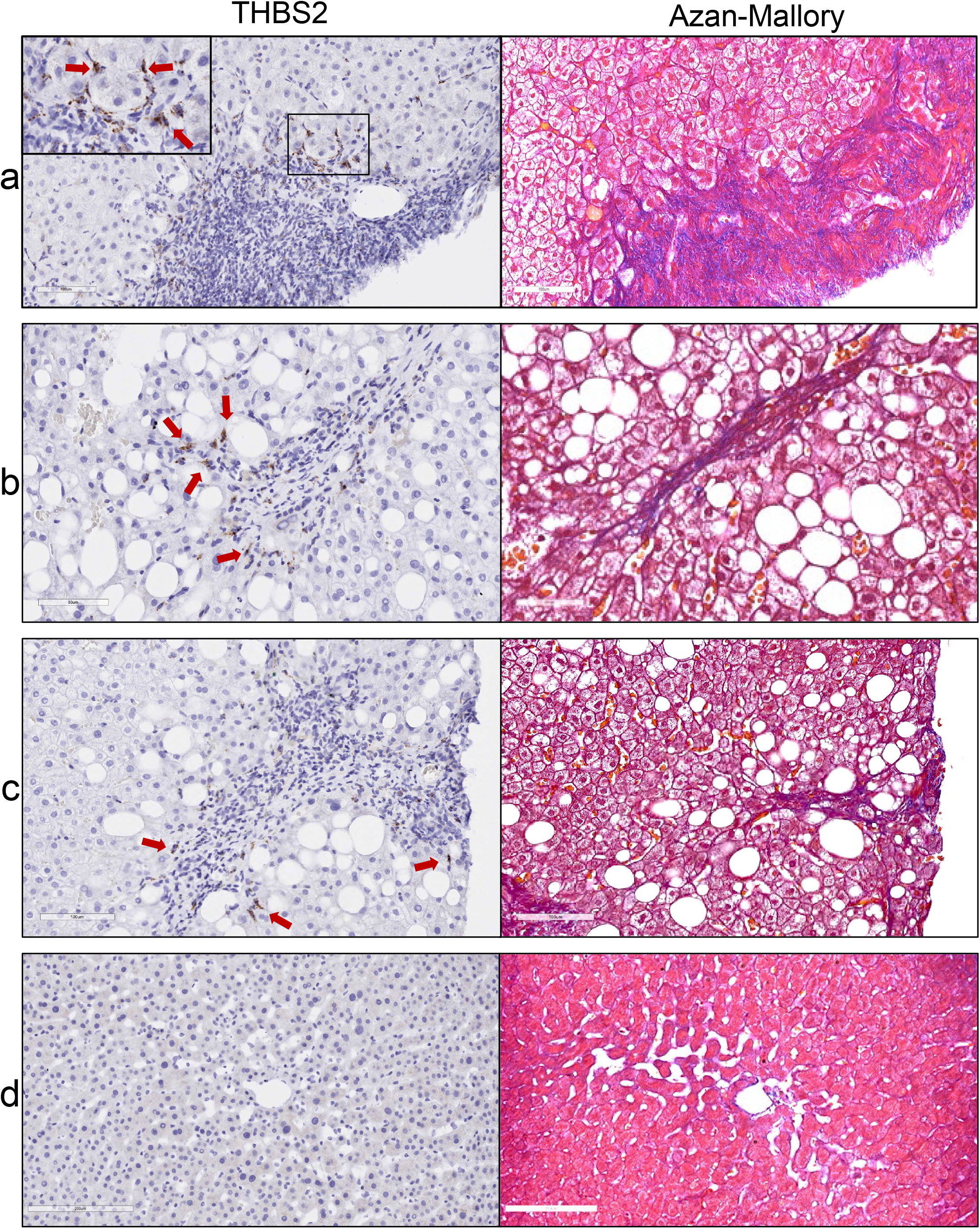
In situ hybridization analysis of NAFLD liver reveals THBS2 expression in HSCs around collagen fibers. **a-d,** In situ hybridization analysis of liver tissue samples using a THBS2 mRNA probe (left) and Azan-Mallory staining (right). NAFLD cases with fibrosis (F3, n=3) (**a-c**) and control case without chronic liver disease (n=1) (**d**). Arrows indicate representative positive sites.

### THBS2 and COL1A1 are upregulated in LX-2 cells upon TGFβ treatment

To investigate the mechanisms underlying the association between THBS2 and fibrosis development, in vitro studies were performed using LX-2 cells as immortalized HSCs (*26*) based on our finding that HSCs were the primary cells responsible for THBS2 in fibrotic NAFLD cases. COL1A1 mRNA levels were significantly increased by co-treatment with TGFβ in LX-2 cells (P<0.0001), although HepG2 was not (Figure 5a). Since COL1A1 mRNA increased according to TGFβ concentration (Figure 5b), a TGFβ concentration of 4 ng/mL was used throughout this study.

**Fig. 5.**
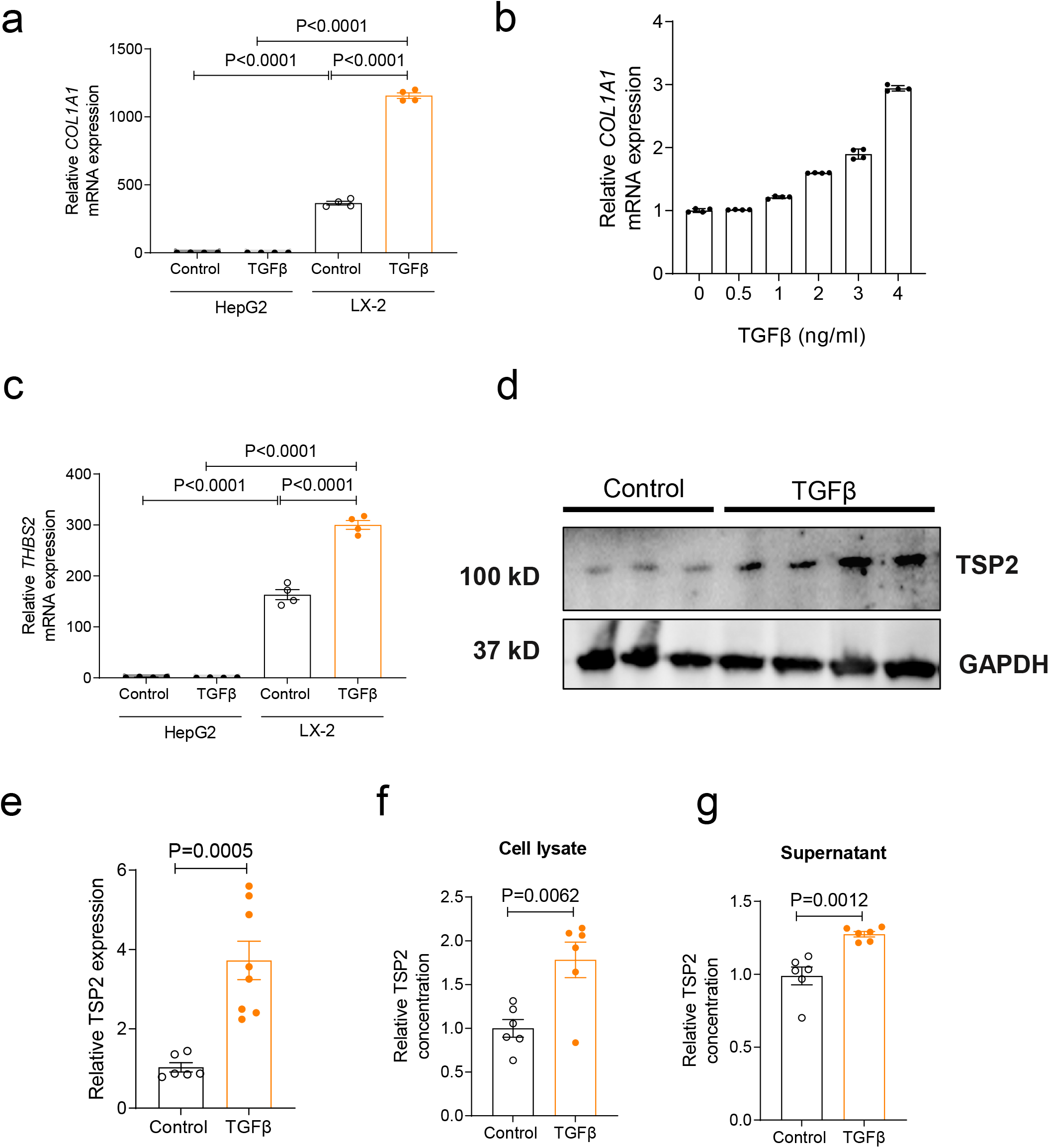
TGFβ-mediated activation of LX-2 cells further raises expression of THBS2/TSP2 and COL1A1. **a,** COL1A1 mRNA expression levels in HepG2 and LX-2 cells in the presence and absence (control) of TGFβ. **b,** COL1A1 mRNA expression levels at different TGFβ concentrations. **c,** THBS2 mRNA expression levels in HepG2 and LX-2 cells in the presence and absence (control) of TGFβ. **d,** Western blotting studies with LX-2 cells treated in the presence and absence (control) of TGFβ. **e-g,** Quantitative analysis of TSP2 protein expression levels in cell lysates by western blotting **(e)** and ELISA **(f),** as well as in the supernatant **(g)** by ELISA of LX-2 cells in the presence and absence (control) of TGFβ **a, c,** Data were expressed relative to basal levels measured in the absence of TGFβ with HepG2 cells. **b,** Data were expressed relative to basal levels measured in the absence of TGFβ. **e-g,** TSP2 protein expression levels were normalized relative to protein expression levels of the control. TGFβ treatment was 4 ng/mL for 10 hours, except for **b.** Data are presented as the mean ± standard error of the mean of at least three independent experiments. Correlation analysis was conducted by one-way ANOVA followed by Bonferroni’s post-hoc test **(a, c)** and the two-tailed Student’s *t*-test **(e-g)**.

LX-2 cells expressed greater than 160-fold THBS2 mRNA levels compared with HepG2 cells (Figure 5c). TGFβ-treated LX-2 cells showed significantly higher THBS2 mRNA expression levels than in control conditions (P<0.0001) (Figure 5c). Furthermore, TGFβ treatment increased TSP2 protein levels in LX-2 cells (Figure 5d-f) and LX-2 cell culture medium (Figure 5g). These results indicated that THBS2/TSP2 expression was enhanced in HSCs and TSP2 was secreted by TGFβ treatment (*16*).

### Knockdown of THBS2 gene suppresses COL1A1 expression in LX-2 cells

To evaluate whether COL1A1 was regulated by THBS2, THBS2 gene knockdown was evaluated in LX-2 cells using siRNA technology. As shown in Figure 6a and b, THBS2-siRNA treatment reduced the expression of THBS2 mRNA and TSP2 protein by roughly half versus control siRNA treatment (P=00003). THBS2 gene knockdown suppressed constitutive COL1A1 mRNA levels (P=0.0046) (Figure 6c), which was also observed in TGFβ-treated conditions (P=0.005) (Figure 6d). Conversely, treatment of LX-2 with recombinant TSP2 increased COL1A1 mRNA levels relative to control conditions (P=0.0051) (Figure 6e).

**Fig. 6.**
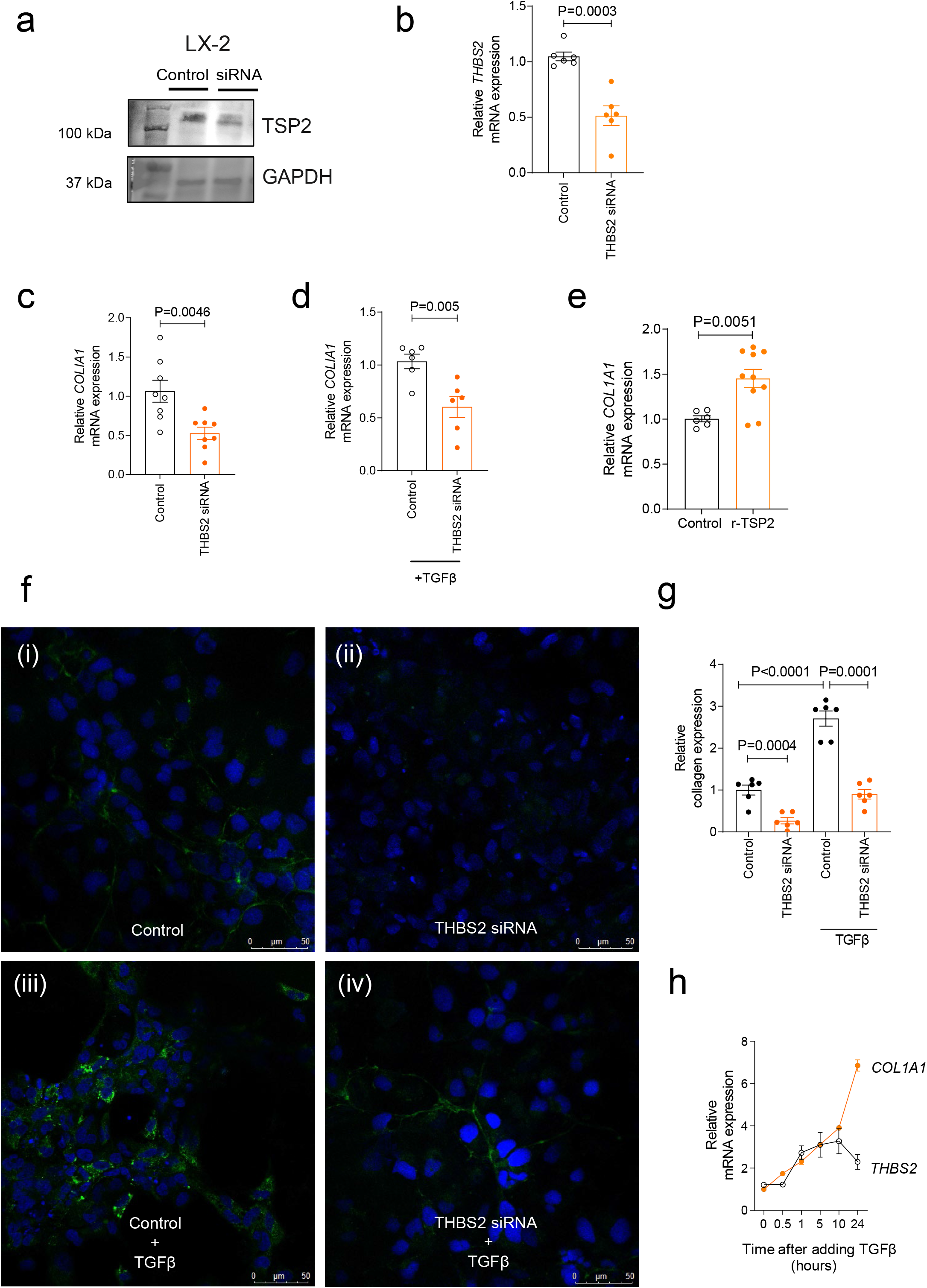
Knockdown of THBS2 gene decreases collagen formation in LX-2 cells. **a, b,** Western blotting studies **(a)** and relative THBS mRNA expression levels **(b)** in LX-2 cells treated with control siRNA and THBS2 siRNA. **c, d,** Relative COL1A1 mRNA expression levels treated with control siRNA and THBS2 siRNA in the presence **(d)** and absence **(c)** of TGFβ. **e,** Relative COL1A2 mRNA expression levels in the presence and absence of recombinant TSP2 protein (30 ng/mL). **f,** Fluorescent staining of LX-2 cells treated with control siRNA or THBS2 siRNA or in the presence and absence of TGFβ: (i) control siRNA, (ii) THBS2 siRNA, (iii) control siRNA and TGFβ, and (iv) THBS2 siRNA and TGFβ. Collagen expression was identified as green and cell nuclei as blue DAPI. **g,** Relative quantification of collagen expression values per cell. **h,** Expression levels of THBS2 mRNA and COL1A1 mRNA over time after addition of TGFβ to LX-2 cells. **b-e, g,** Data were normalized relative to mRNA expression levels of the control. TGFβ treatment was performed with 4 ng/mL for 10 hours, except for **h.** Data are presented as the mean ± standard error of the mean of at least three independent experiments. Correlation analysis was conducted by the two-tailed Student’s *t*-test **(b-e)** and one-way ANOVA followed by Bonferroni’s post-hoc test **(g)**.

Fluorescent staining of type 1 collagen was performed to verify the above results. Green collagen fibers were significantly reduced by THBS2-siRNA (P=0.0004) (Figure 8f, g). TGFβ treatment resulted in prominent collagen accumulation inside and outside of cells, whereas THBS2-siRNA treatment significantly decreased collagen (P=0.0001) (Figure 8f, g). These findings strongly implicated THBS2 as a regulatory gene for collagen formation in LX-2 cells. Examining the time course of TGFβ treatment in LX-2 cells and the expression of THBS2 and COL1A1, THBS2 mRNA expression peaked at 10 hours, followed next by an increase in COL1A1 (Figure 6h). This result was consistent with the notion of THBS2 being a regulator of COL1A1 in LX-2 cells.

### THBS2/TSP2 expression is independent of the classical TGFβ-SMAD pathway in LX-2 cells

It is well known that TGFβ expresses collagen via the phosphorylation of SMAD2/3 (*27*), and so we investigated the association between THBS2/TSP2 expression and SMAD in LX-2 cells. As shown in Figure 7a and b, the phosphorylation of SMAD2/3 induced by TGFβ treatment in LX-2 cells was not affected by the presence or absence of THBS2-siRNA. By examining the changes in THBS2 and COL1A1 mRNA expression levels by SMAD inhibitor treatment in LX-2 cells, we observed that while COL1A1 mRNA levels were significantly reduced (P=0.0005) by the SMAD inhibitor, THBS2 levels were not (Figure 7c). These findings indicated that THBS2/TSP2 expression in LX-2 cells was independent of the classical TGFβ-SMAD pathway (Figure 7d).

**Fig. 7.**
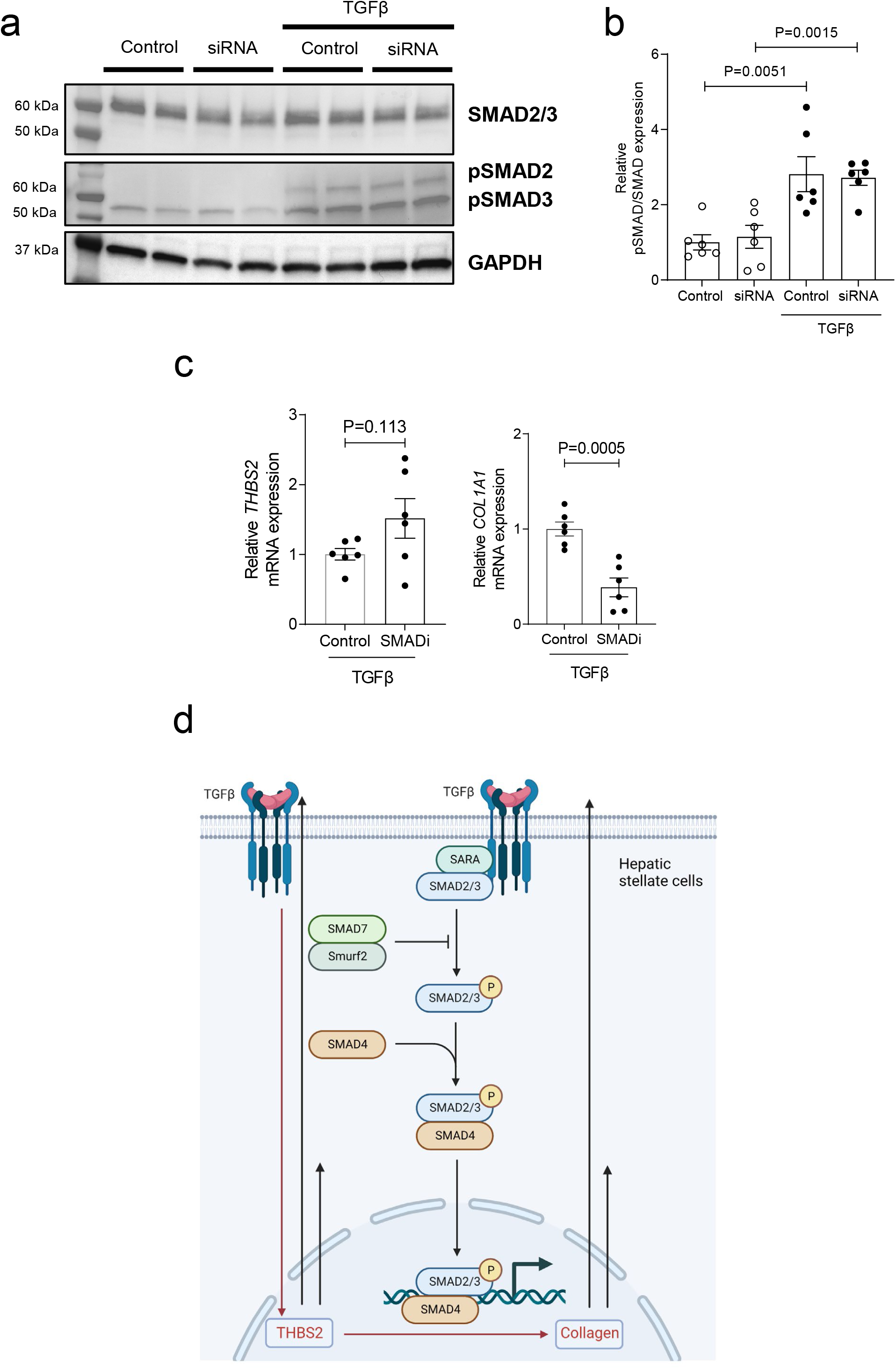
THBS2 expression is independent of TGFβ-SMAD2/3 pathway in LX-2 cells. **a, b,** Western blotting studies of SMAD 2/3 **(a)** and quantitative analysis **(b)** in LX-2 cells treated with control siRNA and THBS2 siRNA in the presence and absence of TGFβ. Control: control siRNA, siRNA: THBS2 siRNA. **c, d,** SMAD inhibitors downregulate COL1A1 mRNA expression levels without affecting THBS2 mRNA expression levels. SMADi: SMAD inhibitor, SIS3 (10 µM). **d,** Proposed mechanism of TGFβ-THBS2/TSP2 expression in LX-2 cells. **b, c,** Data were normalized relative to expression levels of the control. TGFβ treatment was performed with 4 ng/mL for 10 hours. Data are presented as the mean ± standard error of the mean of at least three independent experiments. Correlation analysis was conducted by one-way ANOVA followed by Bonferroni’s post-hoc test **(b)** and the two-tailed Student’s *t*-test **(c)**.

## Discussion

In this study, a comprehensive hepatic gene dataset and in-house biopsy tissue from NAFLD patients revealed significantly higher expression of THBS2 in the livers of patients with advanced fibrosis than in those with mild fibrosis. This result is consistent with several previous reports and highlights the significance of THBS2 in NAFLD fibrosis (*15, 16*). Interestingly, detailed gene network analysis and qPCR examination showed that the THBS2 gene was strongly associated with COL1A1, COL1A2, COL3A1, and COL4A1 expression, which supported the relationship between THBS2 and collagen expression witnessed in cell experiments.

In the analysis of scRNA-seq data, a greater number of THBS2-expressing cells were observed in the HSC cluster. Additionally, the THBS2 expression level of HSCs in NAFLD cases with fibrosis was higher than that of HSCs in NAFLD/healthy individuals without fibrosis. This was consistent with a very recent scRNA-seq data analysis study showing THBS2 expression to be more abundant in fibroblasts in the liver (*28*). To confirm this notion, we performed in situ hybridization analysis of liver tissue from NAFLD patients with severe fibrosis to reveal high THBS2 expression in HSCs at the margins of collagen fiber accumulation. These results prompted us to hypothesize that HSCs played a certain role in collagen formation; indeed, TGFβ treatment markedly increased THBS2/TSP2 expression and simultaneously promoted collagen formation in experiments with LX-2 cells. The suppression of THBS2 reduced collagen expression, identifying it as a possible upstream regulator of collagen expression.

ECM is composed of numerous proteins, including collagen, fibronectin, and laminin (*29*). Genetic mutations of ECM proteins can profoundly alter ECM properties, as evidenced by Marfan and Ehlers-Danlos syndromes (*14*). On the other hand, TSP2 does not directly contribute to ECM structure or stability, but may influence cell-matrix interactions and cell signaling/behavior (*30*). For instance, although whole-body TSP2-deficient mice had a normal appearance and were capable of normal reproduction, the animals had loosened connective tissue, including the skin, ligaments, and tendons, which appeared as disorganized and abnormal collagen fibers on microscopy (*31*). Such results emphasize that TSP2 is required for proper formation and organization of collagen fibers in the skin and tendons. It is reasonable to presume that TSP2 can also affect collagenogenesis in the liver in a similar manner.

Little is known on the transcriptional regulation of TSP2 (*14*). However, the regulatory mechanism of thrombospondin 1 (TSP1), which shares 85% of amino acids along with structural similarities and a similar binding domain, may provide a stepping stone to understand the regulatory mechanism of TSP2 (*14*). The binding domain of TSP1 is postulated to interact with cell surface receptors (LRP, CD36, and CD47), ECM components (decorin, fibronectin, and heparan sulfate proteoglycan), enzymes (matrix metalloproteinase, elastase, and cathepsin G), and calcium as well as TGFβ (*14*). The interaction of TGFβ with this binding domain is consistent with the increased expression of THBS2/TSP2 upon TGFβ loading in LX-2 cells in the present study.

Lastly, TGFβ is a multifunctional cytokine that plays an important role in regulating cell proliferation, differentiation, migration, and ECM deposition (*32*). TGFβ signaling is essential for the formation and maintenance of tissue homeostasis, and its dysregulation is strongly implicated in NAFLD fibrosis (*33, 34*). In the liver, TGFβ is produced primarily by Kupffer cells and hepatic sinusoidal endothelial cells (*35*). During liver injury, TGFβ promotes the activation and proliferation of HSCs as well as the formation of ECM components, especially collagen (*36, 37*). The collagen formation in HSCs is mainly regulated by the TGFβ-SMAD2/3 pathway (*38*). However, in the present experiments, whereas TGFβ-induced collagen synthesis was inhibited by disrupting THBS2/TSP2 expression, SMAD2/3 phosphorylation was unaltered in LX-2 cells. The detailed regulation of collagen by TSP2 is complex and may involve a variety of regulatory systems, and thus is a subject for future studies.

In summary, this investigation revealed that HSCs express THBS2/TSP2 in the liver of NAFLD patients with advanced fibrosis using in situ hybridization and a scRNA-seq dataset. We also provided evidence that THBS2 knockdown inhibited collagen formation in the LX-2 cell line of HSCs. These results suggest the potential clinical utility of THBS2 as a therapeutic target to halt the progression of liver fibrosis in NAFLD.

## Materials and Methods

### Liver mRNA data collection and processing

The mRNA expression data from the Gene Expression Omnibus (GEO) database was downloaded through a microarray dataset (GSE49541) for processing by GEO2R and incorporation into this study (*21, 22*). The data compared the expression of various human genes at different liver fibrosis stages in NAFLD patients. Differentially expressed genes were screened from the GEO dataset with a threshold of |log2FC|>1 and adjusted P<0.02 (*19*).

### Pathway enrichment analysis

The GSE49541 dataset was subjected to enrichment and network analysis using Metascape (https://Metascape.org/) (*21–23*). Metascape is a web-based portal designed to provide a comprehensive gene list annotation and analysis resource for biologists (*23*). To gain insights into the biological roles of identified differentially expressed genes (DEGs) and differentially expressed proteins (DEPs), we conducted pathway enrichment analysis of Gene Ontology Biological Process, Kyoto Encyclopedia of Genes and Genomes, Reactome, and Canonical pathway in Metascape tools (*23, 39–41*). By inputting the lists of DEGs and DEPs simultaneously, Metascape can identify commonly enriched and selectively enriched pathways from two levels, which enables a comprehensive assessment of the molecular features of the biological process.

### scRNA-seq dataset processing

In this study, the scRNA-seq datasets GSE174748 and GSE189175 (*24, 25*) were analyzed using the Seurat R toolkit 4.5. Initially, quality control and filtering were performed to remove lower quality cells (>10% mitochondrial genes), followed next by normalization and scaling. Based on the information provided, condition, sex, and tissue type were annotated in the metadata, which was later used in downstream analysis. After quality control & normalization, the datasets were merged using the canonical correlation analysis function and subsequently integrated using harmony to remove any batch effects. Upon integration, the final dataset was subjected to normalization and scaling once again. The Top 2000 variable feature was employed using Seurat’s built-in function to construct principal components, with the top 15 principal components used for clustering and manifold approximation and projection for dimension reduction visualization. Clusters were annotated using known canonical cell-specific markers.

### Liver samples from NAFLD patients and histological findings

This study included liver tissue samples from 96 biopsy-proven Japanese NAFLD patients admitted to Shinshu University Hospital (Matsumoto, Japan) between 2015 and 2018. The clinical data of the patients are summarized in Table 1. This study was reviewed and approved by the Institutional Review Board of Shinshu University Hospital (Matsumoto, Japan) (approval number: 3021), and written informed consent was obtained from all participating subjects. The investigation was conducted according to the principles of the Declaration of Helsinki. Liver specimens of at least 1.5 cm in length were obtained from segments 5 or 8 using a 14-gauge needle as described previously and immediately fixed in 10% neutral formalin (*42*). Sections of 4 μm in thickness were cut and stained using the hematoxylin and eosin and Azan-Mallory methods. The histological activity of NAFLD was assessed by an independent expert pathologist in a blinded manner according to the NAFLD scoring system (*43*). Fibrosis stage was scored as follows: F0, none; F1, perisinusoidal or periportal; F2, perisinusoidal and portal/periportal; F3, bridging fibrosis; and F4, cirrhosis (*43*).

### THBS2 RNA in situ hybridization

Detection of THBS2 mRNA was performed using an RNAscope kit (Cosmo Bio), according to the manufacturer’s instructions using unstained tissue sections as described in a previous report (*44*). Brown punctate dots in the nucleus and/or cytoplasm indicated positive staining (*44*).

### Quantitative reverse-transcription polymerase chain reaction analysis of gene expression

Total RNA was extracted from frozen tissues or cultured cells using the RNeasy mini kit (Qiagen). SuperScript III First-Strand Synthesis SuperMix (Invitrogen) was used to prepare cDNA. The SYBR green method (Applied Biosystems) was employed for quantitative polymerase chain reaction (qPCR) studies (*45*). ΔΔCt was used to normalize the gene expression data collected in this study relative to the expression of GAPDH (*46*). The PCR primers used in this study are listed in Table 2.

**Table 2.**
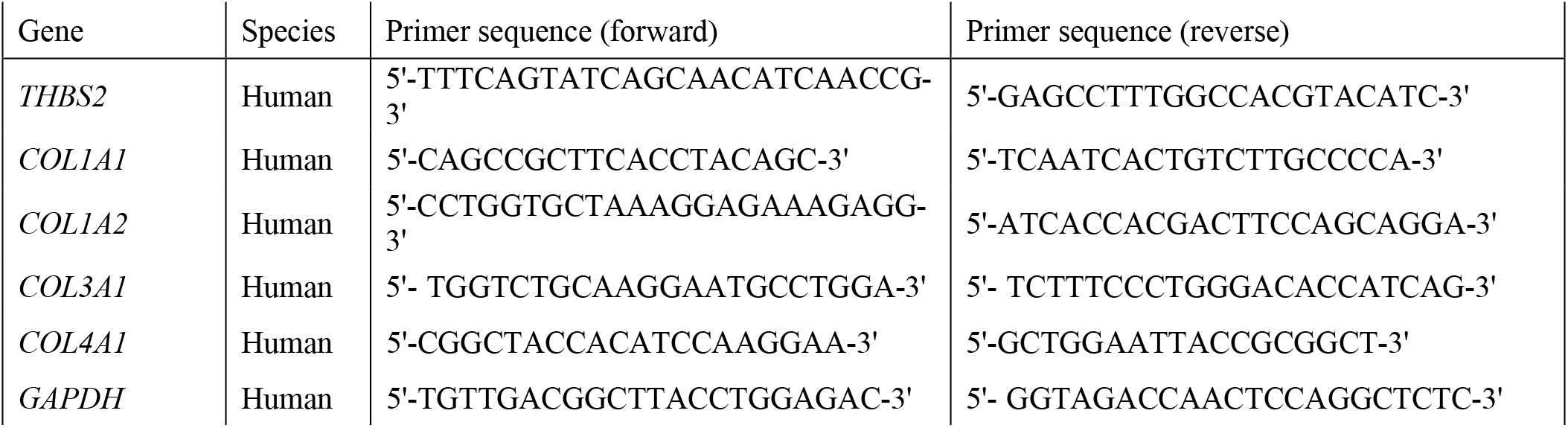
Primers used for qPCR studies.

### Western blotting studies

LX-2 cells were lysed and proteins were extracted using radioimmunoprecipitation assay buffer supplemented with complete EDTA-free protease inhibitor cocktail (Sigma-Aldrich). After centrifugation of cell lysates at 12,000 x g for 10 min, protein concentrations were determined by means of a BCA protein assay kit (Pierce). Subsequently, protein samples were denatured at 95°C using NuPAGE LDS sample buffer (Thermo Fisher Scientific) and separated by 4-12% sodium dodecyl sulfate-polyacrylamide gel electrophoresis. The proteins were then transferred to nitrocellulose membranes and incubated overnight at 4°C with primary antibodies. On the next day, the membranes were washed thoroughly, and then incubated with horseradish peroxidase-conjugated anti-rabbit or anti-goat secondary antibodies, followed by visualization of the separated protein bands using SuperSignal West Pico Chemiluminescent Substrate (Thermo Fisher Scientific) on a c600 Imaging System Imager (Azure Biosystems) (*47*). Immunoreactive bands were quantified using Image J Software (NIH) (*48*).

The following antibodies (source, catalog #, and dilution are indicated) were used: TSP2 (Abcam, #ab112543, 1:1000), small mothers against decapentaplegic (SMAD) 2/3 (D7/G7) (Cell Signaling Technology, #8685, 1:1,000), Phospho-Smad2 (Ser465/467)/Smad3 (Ser423/425) (D27/F4) (Cell Signaling Technology, #8828, 1:1,000), GAPDH (D16/H11) (Cell Signaling Technology, #5174, 1:1,000), anti-rabbit IgG, and HRP-linked antibody (Cell Signaling Technology #7074, 1:2,000).

### Cells

LX-2 cells were obtained from Sigma-Aldrich and HepG2 cells were procured from Cellular Engineering Technologies. Both cell lines were cultured in Dulbecco’s Modified Eagle Medium with 2% fetal bovine serum (*49, 50*).

### siRNA-mediated knockdown of THBS2 expression in LX-2 cells

Human THBS2 siRNA and control siRNA were purchased from Santa Cruz. On day 0, LX-2 cells were transfected with siRNA using Lipofectamine RNAiMAX (Thermo Fisher Scientific) according to the manufacturer’s instructions (*48*). Four hours later, the cells were transferred to normal medium and used for further experiments on day 3.

### Immunocytochemistry

LX-2 cells were plated on glass coverslips (Thermo Fisher Scientific) in 12-well culture dishes and grown to approximately 50% confluence for three days to promote cell adherence. The cells were then washed twice with cold serum-free medium, and then fixed in 4% paraformaldehyde in phosphate-buffered saline (PBS) for 10 minutes. After fixation, the cells were washed twice with PBS followed by permeabilization with PBS containing 0.1% Triton X-100 for 15 minutes. The cells were next washed twice with PBS and incubated with blocking solution (5% BSA in PBS) for 30 minutes. Primary antibodies (collagen type I obtained from Cosmo Bio, diluted 1:100 in blocking solution) were incubated with the cells for one hour. After three washes with 0.2% Tween 20 in PBS (PBST), the cells were incubated with fluorescein-labelled anti-mouse IgG (Vector Laboratories, diluted 1:200 in blocking buffer) for one hour. The cells were washed three times with PBST, stained with DAPI (4′,6-diamidino-2-phenylindole molecular probes, diluted 1:5000 in PBS) for one minute, and then washed three times with PBST and twice with PBS. The cells were viewed with a Nikon Eclipse E600 fluorescence microscope (*26*). Image J technology (NIH) was employed for the quantification of collagen (*51*).

### Compounds and enzyme-linked immunosorbent assays

The recombinant human transforming growth factor beta (TGFβ) protein and recombinant human TSP2 protein were obtained from R&D Systems. The SMAD3 Inhibitor SIS3 was procured from Cayman Chemical Company. Compound concentrations and treatment times are shown in each Figure legend. TSP2 concentrations in LX-2 cell lysates were determined using enzyme-linked immunosorbent assays following the manufacturer’s instructions (Quantikine® ELISA, #DTSP20, R&D Systems) (*16*). The cell lysate method for LX-2 cells was identical to that for western blotting.

### Statistics

Data were collected and analyzed using Prism 8 (GraphPad) and StatFlex Ver. 7.0. All data are expressed as the mean ± standard error of the mean for the indicated number of observations. Prior to the specific statistical tests, we performed testing for normality and homogeneity of variance. The data were then evaluated for statistical significance by one-way ANOVA followed by the indicated post hoc test or the two-tailed unpaired Student’s *t*-test, as appropriate. Correlation analysis was conducted by Spearman’s test. A P value of <0.05 was considered statistically significant.

### Data availability

The gene data referenced during the study have been deposited in GEO under the accession codes GSE49541, GSE174748, and GSE189175. All the other data supporting the findings of this study are available within the article and its Supplementary Information files or from the corresponding author on reasonable request.

## Acknowledgments

This research was supported by AMED under grant number JP23fk0210125 and by JSPS KAKENHI grant number JP22K20884. T.K. was the recipient of a the Aiba Works Medical Research Grant for carrying out several experiments. The authors thank Asami Yamazaki and Trevor Ralph for their assistance in sample preparation and English proofreading, respectively.

## Author contributions

T.K., T.I., and N.T. designed the study and researched data. T.K., T.I., T.Y., T.N., and D.A. carried out experiments and interpreted and analyzed experimental data. Y.Y. and S.J. collected patient samples and analyzed data. S.W., S.K., H.Z., and S.P.P. analyzed scRNA-seq data. M.I. and T.U. provided pathological guidance. T.K. and T.I. wrote the manuscript. S.P.P. and T.U. supervised the entire study.

---

**Supp. Fig. 1.**
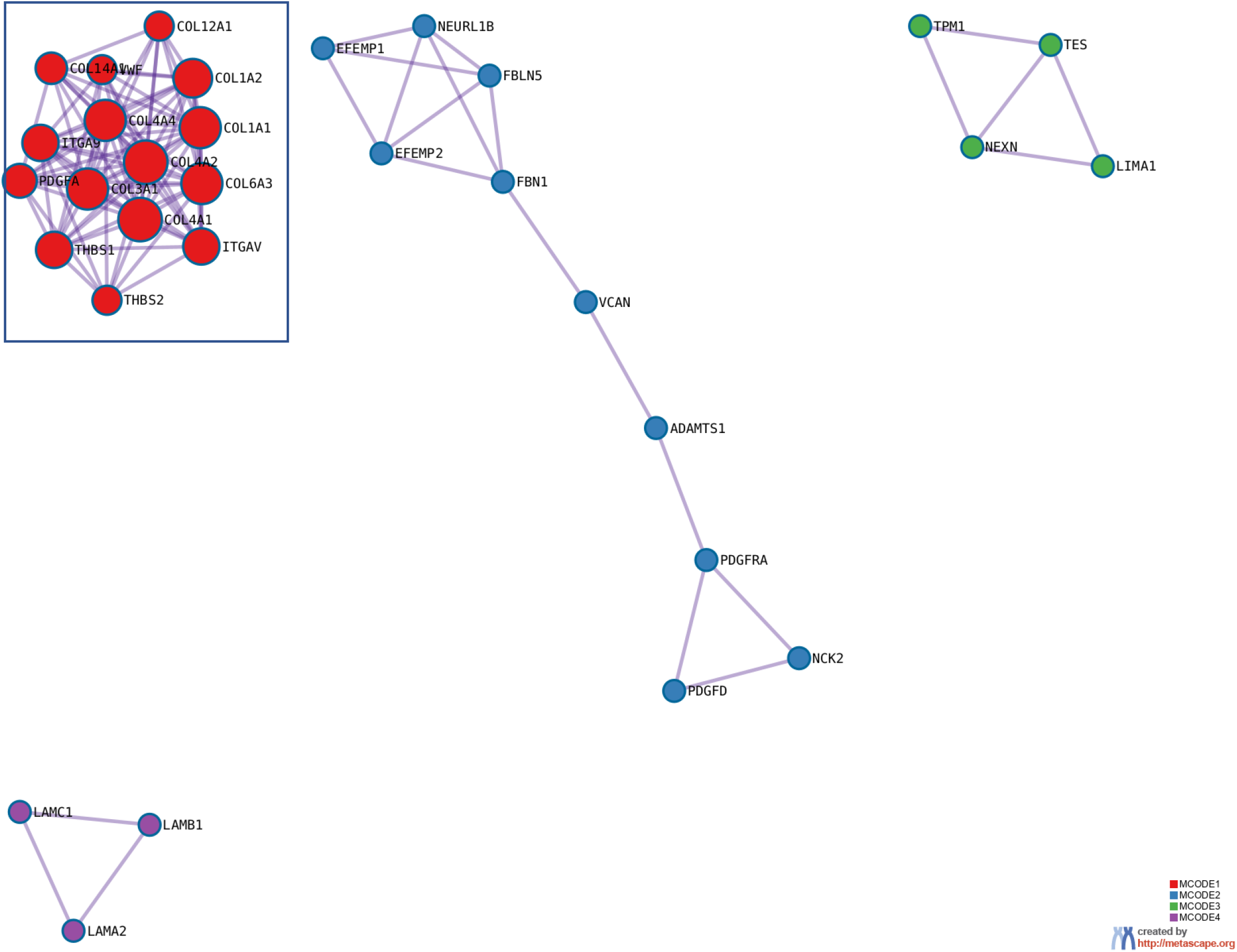
Network analysis of NAFLD liver gene set (GSE49541) using Metascape.

## References

1. J. V. Lazarus et al., Advancing the global public health agenda for NAFLD: a consensus statement. Nat Rev Gastroenterol Hepatol 19, 60–78 (2022).

2. E. E. Powell, V. W. Wong, M. Rinella, Non-alcoholic fatty liver disease. Lancet 397, 2212–2224 (2021).

3. Z. Younossi et al., Global burden of NAFLD and NASH: trends, predictions, risk factors and prevention. Nat Rev Gastroenterol Hepatol 15, 11–20 (2018).

4. N. Tanaka et al., Current status, problems, and perspectives of non-alcoholic fatty liver disease research. World J Gastroenterol 25, 163–177 (2019).

5. E. Vilar-Gomez et al., Fibrosis Severity as a Determinant of Cause-Specific Mortality in Patients With Advanced Nonalcoholic Fatty Liver Disease: A Multi-National Cohort Study. Gastroenterology 155, 443–457 e417 (2018).

6. T. Kimura, N. Tanaka, E. Tanaka, What will happen in patients with advanced nonalcoholic fatty liver disease? Hepatobiliary Surg Nutr 8, 283–285 (2019).

7. A. J. Sanyal et al., Prospective Study of Outcomes in Adults with Nonalcoholic Fatty Liver Disease. N Engl J Med 385, 1559–1569 (2021).

8. K. Tokushige et al., Evidence-based clinical practice guidelines for nonalcoholic fatty liver disease/nonalcoholic steatohepatitis 2020. J Gastroenterol 56, 951–963 (2021).

9. T. Kimura, S. Singh, N. Tanaka, T. Umemura, Role of G Protein-Coupled Receptors in Hepatic Stellate Cells and Approaches to Anti-Fibrotic Treatment of Non-Alcoholic Fatty Liver Disease. Front Endocrinol (Lausanne) 12, 773432 (2021).

10. K. Zhang, M. Li, L. Yin, G. Fu, Z. Liu, Role of thrombospondin-1 and thrombospondin-2 in cardiovascular diseases (Review). Int J Mol Med 45, 1275–1293 (2020).

11. F. Gao et al., Diagnostic and Prognostic Roles of Thrombospondin-2 in Digestive System Cancers. Biomed Res Int 2022, 3749306 (2022).

12. R. Simantov, M. Febbraio, R. L. Silverstein, The antiangiogenic effect of thrombospondin-2 is mediated by CD36 and modulated by histidine-rich glycoprotein. Matrix Biol 24, 27–34 (2005).

13. M. Rienks, A. P. Papageorgiou, N. G. Frangogiannis, S. Heymans, Myocardial extracellular matrix: an ever-changing and diverse entity. Circ Res 114, 872–888 (2014).

14. N. E. Calabro, N. J. Kristofik, T. R. Kyriakides, Thrombospondin-2 and extracellular matrix assembly. Biochim Biophys Acta 1840, 2396–2402 (2014).

15. K. Kozumi et al., Transcriptomics Identify Thrombospondin-2 as a Biomarker for NASH and Advanced Liver Fibrosis. Hepatology 74, 2452–2466 (2021).

16. T. Kimura et al., Serum thrombospondin 2 is a novel predictor for the severity in the patients with NAFLD. Liver Int 41, 505–514 (2021).

17. C. H. Lee et al., Circulating Thrombospondin-2 as a Novel Fibrosis Biomarker of Nonalcoholic Fatty Liver Disease in Type 2 Diabetes. Diabetes Care 44, 2089–2097 (2021).

18. X. Wu et al., Serum Thrombospondin-2 Levels Are Closely Associated With the Severity of Metabolic Syndrome and Metabolic Associated Fatty Liver Disease. J Clin Endocrinol Metab 107, e3230–e3240 (2022).

19. T. Iwadare et al., Circulating thrombospondin 2 levels reflect fibrosis severity and disease activity in HCV-infected patients. Sci Rep 12, 18900 (2022).

20. T. Matsumae et al., Thrombospondin-2 as a Predictive Biomarker for Hepatocellular Carcinoma after Hepatitis C Virus Elimination by Direct-Acting Antiviral. Cancers (Basel) 15, (2023).

21. C. A. Moylan et al., Hepatic gene expression profiles differentiate presymptomatic patients with mild versus severe nonalcoholic fatty liver disease. Hepatology 59, 471–482 (2014).

22. S. K. Murphy et al., Relationship between methylome and transcriptome in patients with nonalcoholic fatty liver disease. Gastroenterology 145, 1076–1087 (2013).

23. Y. Zhou, et al., Metascape provides a biologist-oriented resource for the analysis of systems-level datasets. Nat Commun 10, 1523 (2019).

24. A. Filliol et al., Opposing roles of hepatic stellate cell subpopulations in hepatocarcinogenesis. Nature 610, 356–365 (2022).

25. M. Alvarez et al., Human liver single nucleus and single cell RNA sequencing identify a hepatocellular carcinoma-associated cell-type affecting survival. Genome Med 14, 50 (2022).

26. L. Xu et al., Human hepatic stellate cell lines, LX-1 and LX-2: new tools for analysis of hepatic fibrosis. Gut 54, 142–151 (2005).

27. H. Khalil et al., Fibroblast-specific TGF-beta-Smad2/3 signaling underlies cardiac fibrosis. J Clin Invest 127, 3770–3783 (2017).

28. O. Govaere et al., A proteo-transcriptomic map of non-alcoholic fatty liver disease signatures. Nat Metab, (2023).

29. J. Halper, M. Kjaer, Basic components of connective tissues and extracellular matrix: elastin, fibrillin, fibulins, fibrinogen, fibronectin, laminin, tenascins and thrombospondins. Adv Exp Med Biol 802, 31–47 (2014).

30. P. Bornstein, E. H. Sage, Matricellular proteins: extracellular modulators of cell function. Curr Opin Cell Biol 14, 608–616 (2002).

31. T. R. Kyriakides et al., Mice that lack thrombospondin 2 display connective tissue abnormalities that are associated with disordered collagen fibrillogenesis, an increased vascular density, and a bleeding diathesis. J Cell Biol 140, 419–430 (1998).

32. N. R. Gough, X. Xiang, L. Mishra, TGF-beta Signaling in Liver, Pancreas, and Gastrointestinal Diseases and Cancer. Gastroenterology 161, 434–452 e415 (2021).

33. H. Ahmed et al., TGF-beta1 signaling can worsen NAFLD with liver fibrosis backdrop. Exp Mol Pathol 124, 104733 (2022).

34. I. Fabregat et al., TGF-beta signalling and liver disease. Febs J 283, 2219–2232 (2016).

35. A. Carambia et al., TGF-beta-dependent induction of CD4(+)CD25(+)Foxp3(+) Tregs by liver sinusoidal endothelial cells. J Hepatol 61, 594–599 (2014).

36. T. Tsuchida, S. L. Friedman, Mechanisms of hepatic stellate cell activation. Nat Rev Gastroenterol Hepatol 14, 397–411 (2017).

37. F. Jia, X. Hu, T. Kimura, N. Tanaka, Impact of Dietary Fat on the Progression of Liver Fibrosis: Lessons from Animal and Cell Studies. Int J Mol Sci 22, (2021).

38. L. R. Ellis, D. R. Warner, R. M. Greene, M. M. Pisano, Interaction of Smads with collagen types I, III, and V. Biochem Biophys Res Commun 310, 1117–1123 (2003).

39. C. Gene Ontology, The Gene Ontology project in 2008. Nucleic Acids Res 36, D440–444 (2008).

40. M. Kanehisa, Y. Sato, M. Kawashima, M. Furumichi, M. Tanabe, KEGG as a reference resource for gene and protein annotation. Nucleic Acids Res 44, D457–462 (2016).

41. B. Jassal et al., The reactome pathway knowledgebase. Nucleic Acids Res 48, D498–D503 (2020).

42. N. Fujimori et al., 2-Step PLT16-AST44 method: Simplified liver fibrosis detection system in patients with non-alcoholic fatty liver disease. Hepatol Res 52, 352–363 (2022).

43. D. E. Kleiner et al., Design and validation of a histological scoring system for nonalcoholic fatty liver disease. Hepatology 41, 1313–1321 (2005).

44. T. Nakajima et al., Distribution of Lgr5-positive cancer cells in intramucosal gastric signet-ring cell carcinoma. Pathol Int 66, 518–523 (2016).

45. F. Jia et al., Dietary Restriction Suppresses Steatosis-Associated Hepatic Tumorigenesis in Hepatitis C Virus Core Gene Transgenic Mice. Liver Cancer 9, 529–548 (2020).

46. S. Jain et al., Chronic activation of a designer G(q)-coupled receptor improves beta cell function. J Clin Invest 123, 1750–1762 (2013).

47. P. Diao et al., Dietary Fat Composition Affects Hepatic Angiogenesis and Lymphangiogenesis in Hepatitis C Virus Core Gene Transgenic Mice. Liver Cancer 12, 57–71 (2023).

48. T. Kimura et al., Adipocyte G(q) signaling is a regulator of glucose and lipid homeostasis in mice.m Nat Commun 13, 1652 (2022).

49. C. Y. Han et al., Hepcidin inhibits Smad3 phosphorylation in hepatic stellate cells by impeding ferroportin-mediated regulation of Akt. Nat Commun 7, 13817 (2016).

50. X. Wu et al., CUG-binding protein 1 regulates HSC activation and liver fibrogenesis. Nat Commun 7, 13498 (2016).

51. T. J. Collins, ImageJ for microscopy. Biotechniques 43, 25–30 (2007).

